# The importance of cellulolytic and xylanolytic activities of *Bacillus subtilis* on dairy cow feed digestibility

**DOI:** 10.1101/2023.03.20.533435

**Authors:** Valeria Bontà, Marco Battelli, Erlinda Rama, Michela Casanova, Lorenzo Pasotti, Gianluca Galassi, Stefania Colombini, Cinzia Calvio

## Abstract

A large body of literature has accumulated on the beneficial impact of the administration of *Bacilli* to dairy cows, particularly on dry matter intake, lactation performances and milk composition. In this work we sought to establish whether the ability of *B. subtilis* to secrete cellulase and xylanase enzymes could be involved in the positive effect exerted by these bacteria.

Several feed ingredients were treated with two *B. subtilis* strains only differing for the amount of secreted cellulosolytic and xylanolytic enzymes, and feed quality was assessed. We found that *in vitro* fibre digestibility correlated with the activity of those enzymes. Our results revealed that *B. subtilis* cellulases and xylanases can effectively improve forage quality, providing a scientific rationale on the use of *Bacilli* as forage supplements to improve animal productivity. Moreover, a particular care was taken in designing a sustainable and economically viable *Bacillus*-based additive preparation process.

## 1. Introduction

Current challenges for the livestock sector are the identification of technologies able to boost animal performances through sustainable farming systems (Capper and Bauman, 2013; Adesogan et al., 2019), without exacerbating the exploitation of natural resources. Sustainability can be met, without expanding land occupation, by raising animal productivity through the improvement of the nutritional properties of current forages or of sub-optimal feeds. Forage fibres are mainly constituted by cellulose and hemicellulose, and animal productivity has been often correlated with their digestibility by the rumen microorganisms (Oba and Allen, 1999; Kendall, 2009). A higher fibre digestibility translates into a lower amount of feed necessary to produce a unit of meat or milk, reducing the impact of animal production on environmental resources. Another non-negligible advantage of incrementing fibre digestibility is the possibility of using less nutritive biomasses that, differently from grains, are not in competition with human nutrition.

To increase fibres surface area accessible to rumen microbial activity, several forage treatments have been considered, ranging from physicochemical approaches, e.g., treatments with strong alkalis, ammonium, urea, or steam, to more environmentally friendly processes based on the application of exogenous hydrolytic enzymes to deconstruct the lignocellulosic biomass (Donnelly et al., 2018; Mor et al., 2018; Adesogan et al., 2019). However, the latter strategy heavily impacts on farm profitability due to the high costs of industrial enzymes (Ferreira et al., 2021); moreover, enzymatic activity is often rapidly lost within the rumen environment (Adesogan et al., 2019). The direct supplementation of microorganisms producing degradative enzyme into forages and diets may overcome such limits, guaranteeing continuous secretion and turnover of hydrolytic enzymes to deconstruct the biomass, facilitating its fermentation in the rumen (Adesogan et al., 2019). Numerous microorganisms are known to secrete large amounts of hydrolytic enzymes that attack the lignocellulosic matrix of vegetable material, mainly represented by fungi and soil bacteria (Himmel et al., 2010). One of the best characterized bacterial species known to produce massive amounts of degradative enzymes is *Bacillus subtilis* (Danilova and Sharipova, 2020). Its attractiveness derives from its propensity to secrete high levels of enzymes directly into the growth medium (up to 20-25 g/litre), the ease of cultivation, and its recognized safety in human and animal nutrition (Schallmey et al., 2004; Westers et al., 2004). Such features, along with its plasticity for genetic engineering that allows to boost enzymatic yields (Van Dijl and Hecker, 2013), the natural safety of its products, and cheap feedstock demand, justify the pivotal position of *B. subtilis* among the industrially relevant microorganisms (Su et al., 2020). Dwelling mainly in the upper layers of soil and in the plant rhizosphere, *B. subtilis* genome has evolutionary accumulated a large set of degradative enzymes (Earl et al., 2008); according to the Carbohydrate Active Enzymes database (CAZy; http://www.cazy.org/), the genome of the common laboratory strain of *B. subtilis* JH642 (Srivatsan et al., 2008; Smith et al., 2014) carries several genes encoding secreted enzymes involved in complex carbohydrates degradation (Lombard et al., 2016). This species, listed in the *Qualified Presumption of Safety (QPS) recommended microorganisms intentionally added to food or feed* (EFSA BIOHAZ Panel, 2022, 2023a, 2023b), was shown to colonize the gastrointestinal tract of mammals and poultry, in the form of vegetative cells (spores) or forming biofilms, thereby acting as probiotic (Bernardeau et al., 2017; Lee et al., 2019; Mun at al., 2021). In particular, *B. subtilis* spores added to dairy cattle feed consistently exerted a positive influence on ruminal fermentation, growth, lactation performance, and milk composition (Sun et al., 2011; Sun et al., 2013; Sun et al., 2016; Souza et al., 2017; Choonkham et al., 2020; Wang et al., 2022; Jia et al., 2022).

Recently, from a common laboratory strain of *B. subtilis*, we derived an isogenic strain in which endogenous cellulases and xylanases production was optimized (Doria et al., 2022). These two strains do not differ for amylase, pullulanase and other hydrolytic enzymes biosynthesis. Several types of common cattle feeds were parallelly treated with the two *B. subtilis* strains, to evaluate whether digestibility of the vegetable fibres, a fundamental parameter in feed quality (Weinberg et al., 2009), could be correlated to the amount of cellulases and xylanases applied to forages.

Besides, to preserve the affordability of the forage treatment, *B. subtilis* was conveniently grown in a salt medium in which the carbon source derived from rice straw, a low-cost agricultural waste, and whole cultures were directly applied to forages, without the need of complex and expensive purification steps.

## 2. Materials and Methods

### 2.1. Bacillus subtilis strains

*Bacillus subtilis* strains used in this study are the wild-type (WT) PB5700 strain and its engineered version PB5703 (PB2OPT). PB5700 is a spontaneous *swrA*^+^ derivative of the JH642 strain (GenBank accession no. CM000489.1) in which the auxotrophies caused by the *trpC2 pheA1* mutations have been cured as described in Doria et al. (2022). PB2OPT was derived from PB5700 by genetic optimization without the introduction of any exogenous DNA sequence, and is similar to OS58 (Doria et al., 2022). A patent application is under preparation for this strain; for this reason, additional details on the engineered strain are herein undisclosed.

### 2.2. Bacterial medium preparation and its characterization

Open-air dried rice straw (*Oryza sativa L.*) was collected from two local farmers in the Pavia area (northwester part of Italy) and stored in burlap sacks. Upon use, straw was minced using a kitchen blender (Moulinex AY45 Easy Power Blender) and dried to constant weight in a 60 °C oven. One litre tap water was added to 25 grams of straw and the suspension was vigorously stirred at room temperature for 2 hours with a magnetic stirrer. Straw was removed through a 1 mm mesh-strainer covered with two sheets of gauze. The washing liquid (WL) was collected and subjected to low-pressure sterilization (5 psi/~0.35 bar), to avoid sugar caramelization, for 20 minutes and then stored at −20 °C.

To quantify the sugars extracted, 300 μL of WL were brought to a final volume of 500 μL with 50 mM Na-Acetate buffer pH 5 and incubated at 50 °C for 64 h with shaking, either alone or in the presence of 22 μL-enzymes cocktail constituted by 1U Cellulase (product n° 22178, Millipore-Sigma), 1U Beta-glucosidase (product n° 49290, Millipore-Sigma), 1U Xylanase (product n° X2629, Millipore-Sigma) and 0.013 U Cellobiohydrolase I (product n° E6412, Millipore-Sigma). A mock reaction was carried out with the enzymatic cocktail in the absence of substrate. After incubation, samples were centrifuged at 16.873 × g at room temperature for 5 min, filtered through sterile 0.2 μm polyethersulfone (PES) filters and stored at −20 °C until HPLC analysis. Twenty-five microlitre samples were injected through an automatic injector and analysed in an LC-2000 HPLC system (Jasco) equipped with a Supelco C-610H 30 cm x 7.8 mm column (59320-U, Sigma Aldrich) and a RID 10A detector (Shimadzu). For chromatography, 0.1 % H_3_PO_4_ was used as mobile phase with 0.5 mL/min flow rate. Column temperature was kept at 30 °C. Chromatograms were analysed by the ChromNAV 2.0 software (Jasco). The column was previously calibrated with standard solutions of glucose, xylose, galactose, mannose, arabinose and cellobiose, serially diluted in mobile phase to prepare calibration curves based on at least three replicates for each sugar. The lower limit of quantification for all sugars was 0.3 g/litre. Xylose, galactose, and mannose showed the same retention time and the same concentration-peak area calibration lines, thereby enabling the precise quantification of their sum. However, since the rice straw content in galactose and mannose is negligible compared with the one of xylose, we assumed that the concentration of co-eluted xylose, galactose and mannose detected was only due to xylose (Belal, 2013; Krishania et al., 2018). Arabinose and cellobiose could not be detected. Background sugars present in the enzymatic cocktail were subtracted from each sample. For control samples with values below the limit of quantification (LOQ) of the calibration curve (0.3 mg/mL), the LOQ/2 value (i.e., 0.15 mg/mL) was used as background (Bergstrand and Karlsson, 2018). Sugar quantification values are the mean of two independent experiments.

### 2.3. Bacillus subtilis cultures

The growth medium was prepared by adding to WL the following components: 7 g/L K_2_HPO_4_; 2 g/L KH_2_PO_4_; 0.5 g/L Na citrate; 0.1 g/L MgSO_4_; 1 g (NH_4_)2SO_4_; 1 g/L asparagine. PB5700 and PB2OPT spores were revitalized over-night on LB plates. Few single colonies were inoculated in 3 mL of Antibiotic Medium 3 (BD-Difco, New Jersey, U.S) and grown at 37 °C with orbital shaking till culture density was appreciable. Pre-cultures were grown in 20 mL of fresh Antibiotic Medium 3 containing with 5 mg/mL glucose and grown at 37 °C over-night. Fermentation was carried out for 24 hours in the WL-based medium at 37 °C with 150 rpm agitation, starting with an optical density at 600 nm (OD_600_) of 0.2. Each fermentation was independently repeated at least three times.

### 2.4. Cellulase and xylanase activity assays

At the end of the incubation, cellulase (endo-1,4-β-glucanase; EC 3.2.1.4) and endo-xylanase (endo-1,4-β-xylanase; EC 3.2.1.8) activities were assayed in the culture supernatants, after acidification, using the CellG5 and XylX6 kits (Megazyme, Wicklow, Ireland) as previously described (Ermoli et al., 2021).

### 2.5. In vitro NDFd analyses

The effect of the enzymes produced by PB5700 and PB2OPT on the degradability of feeds was analysed using an *in vitro* incubation method aimed to evaluate ruminal digestibility of neutral detergent fibre (NDFd).

Ruminal fluid was collected from three rumen-cannulated cows (non-lactating Holstein-Friesian). The donor animals were handled as outlined by the Directive 2010/63/EU on animal welfare for experimental animals and all animal procedures were conducted under the approval of the University of Milan Ethics Committee for animal use and care and with the authorization of the Italian Ministry of Health (authorization n. 904/2016-PR).

The cows were fed a total mixed ratio (TMR) composed by 66% hay and 34% concentrate twice daily to achieve a Dry Matter Intake (DMI) of 8 kg/d. Rumen liquor was collected two hours after the morning feeding, immediately strained through four layers of cheesecloth into a pre-warmed (39 °C) flask, flushed with CO_2_ and used within one hour from sampling.

Three different treatments were analysed: i) the control treatment, constituted by the WL-based growth medium, in the absence of bacteria; ii) the treatment, constituted by PB5700 culture; iii) the treatment, constituted by PB2OPT culture.

The incubations were conducted on 8 different feed ingredients: 2 different meadow hays, 2 different alfalfa hays, 2 types of corn silages, and 2 types of TMR for dairy cows. All feeds were dried at 60 °C for 48 h in a forced-air oven and ground to pass a 1mm Fritsch mill (Fritsch Pulverisette, Idar-Oberstein, Germany). All the experiments (three treatments x 8 feed ingredients) were independently repeated three times (three incubation runs for each method).

The NDF digestibility was determined at 48 h using the Ankom Daisy II incubator as reported by Spanghero et al. (2019) modified as follows: each jar, containing 250 mg of feed, was inoculated with 665 mL of fresh bacterial culture from one of the three experimental treatments (control, PB5700, PB2OPT), 133 mL of salt mix solution (90 g/L KH_2_PO_4_; 5 g/L NaCl; 5 g/L Urea; 4.5 g/L MgSO_4_*7H_2_0; 1 g/L CaCl_2_*2H_2_0), 532 mL of distilled water, 266 mL of reducing solution (Na_2_CO_3_ 15 g/L; Na_2_S*9H_2_O 1 g/L), and 400 mL of rumen fluid. The NDF content of feed ingredients was determined using the Ankom 200 Fiber Analyzer (Ankom Technology, Macedon, NY, USA) following the procedure reported by Mertens (2002).

### 2.6. Statistical analyses

Data were statistically analysed by the PROC MIXED procedure of SAS 9.4 (SAS Institute Inc., Cary, NC, USA), with the following model:

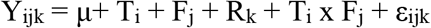

where Y_ijk_ = dependent variable, μ = overall mean, T_i_ = fixed effect of the treatment (i = 1 to 3), F_j_ = fixed effect of the feed (j = 1 to 8), R_k_ = random effect of the fermentation run (k = 1 to 3), T x F_ij_ = interaction treatment x feed, and εijk is the random error. The least-square means were reported. For all statistical analyses, significance was declared at P ≤ 0.05 and trends at P ≤ 0.10.

## 3. Results

### 3.1. B. subtilis feed additive preparation

Supplementation of *Bacillus subtilis* to forages was repeatedly shown to represent a sustainable strategy to increase dairy cows’ performances; however, the acquisition of these additives on the market adds non-negligible costs to farmers. In view of designing an affordable supplement that could be obtained at the farm site, a simplified and inexpensive procedure for the production of *B. subtilis*-based additives was devised. Rice straw is the third most abundant, annually renewable, biomass, and is not in competition with the food or feed industry. Besides a limited use as fodder, there are no value-added applications for this by-product, which is largely burnt on the field, generating environmental problems. To develop a low-cost medium for *B. subtilis*, rice straw represented an attractive feedstock. Water-soluble carbohydrates, i.e., fructose, sucrose and β-1,3-1,4-glucan, are in fact present in significant amounts in rice straw (Park et al., 2009; McIntosh et al., 2011). To recover those sugars through an environmentally friendly and simple process, dried rice straw was vigorously washed with tap water at room temperature. The analysis of the washing liquid (WL) demonstrated the presence of not only monomeric sugars, but also complex sugars, as the amount of the formers increased substantially after enzymatic hydrolysis of WL with commercial enzymes (Table 1).

**Table 1.**
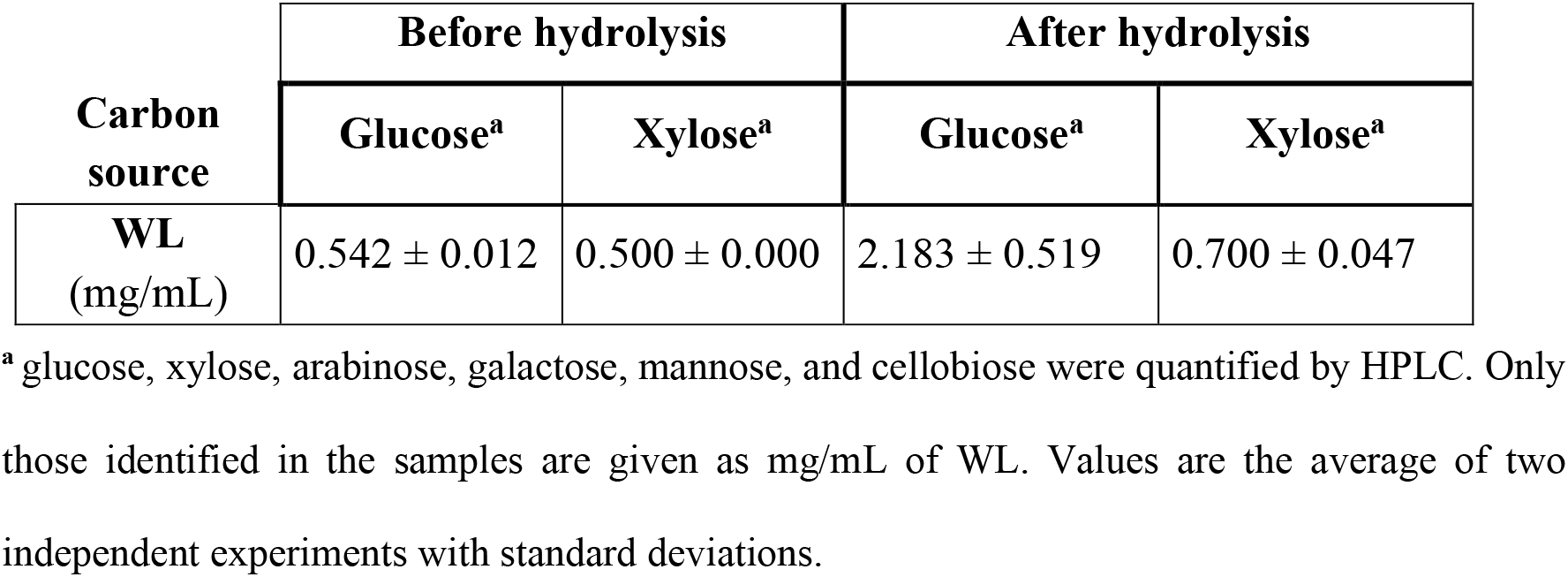
Monomeric sugars contained in WL before and after enzymatic hydrolysis.

The WL fraction was thus used as carbon source for the growth *Bacillus subtilis*. We used the isogenic PB5700 laboratory strain and the optimized PB2OPT strain, just differing for the level of cellulases and xylanases secreted in the growth medium. Both strains were able to grow on the WL fraction, without any preliminary pre-saccharification, reaching a final OD_600_ of 2.85 ± 0.14 for the WT and of 3.83 ± 0.14 for PB2OPT at 24 h. Cellulases and xylanases, released from both strains in the growth medium, were quantified at the end of the incubation. As shown in Fig. 1, secretion of both enzymes was much higher for the optimized strain. Cellulase activity in the supernatant of PB2OPT was 142 folds higher than for PB5700 (92.8 ± 1.6 mU/mL vs 0.7 ± 0.2 mU/mL, respectively). Xylanase activity was also significantly higher in PB2OPT, albeit by only 5.4 folds (18.7 ± 2.2 mU/mL vs 3.5 ± 0.4 mU/mL recorded in the wild-type strain).

**Figure 1.**
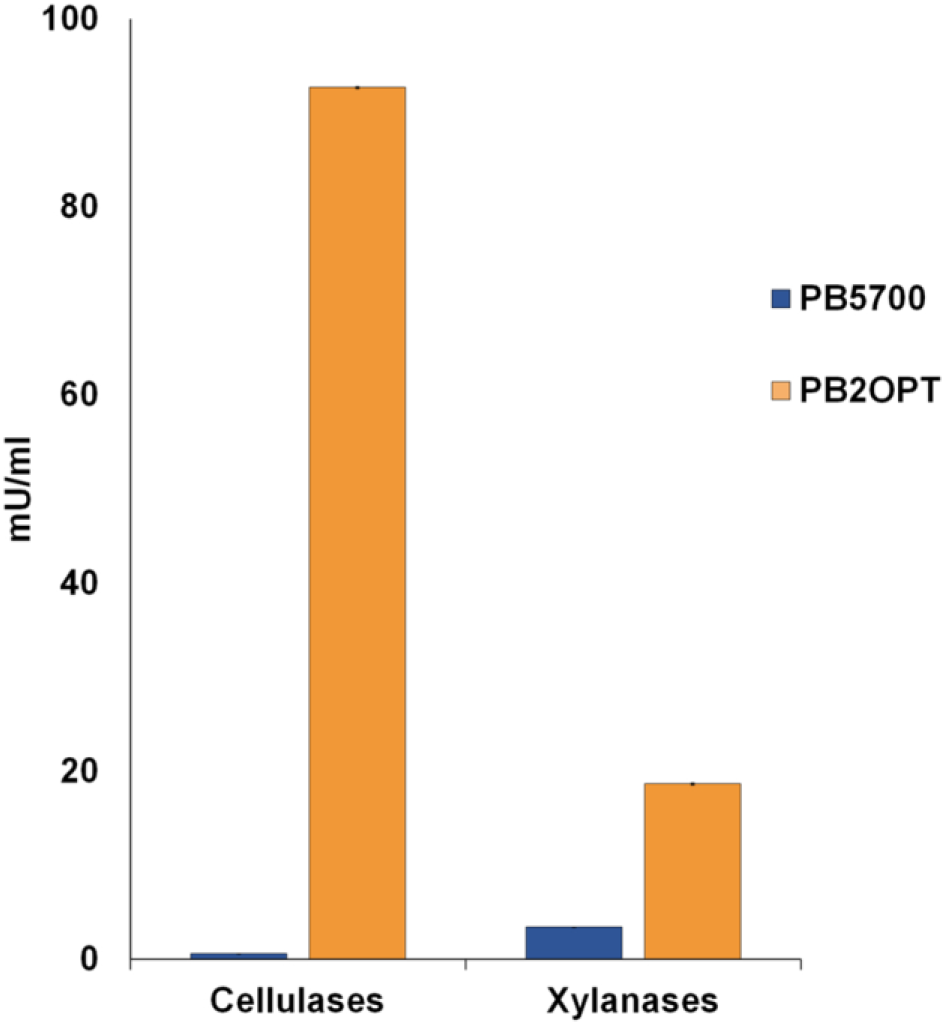
Cellulase and xylanase activities. Cellulase and xylanase activities, expressed in mU per mL of culture media, recorded in the culture supernatant of PB5700 and PB2OPT strains grown on WL medium. Values are the average of at least five independent experiments; error bars (included, although barely visible) represent the standard error of the means.

### 3.2. Feed treatment and fibre digestibility

To preserve the economic affordability of the treatment, raw bacterial cultures were applied to feeds ingredients for dairy cows, such as meadow hay, alfalfa hay, corn silage and TMR, without any preliminary enzyme purification step. Raw cultures of the two *B. subtilis* strains, including bacteria and the growth medium in which degradative enzymes had accumulated, were incubated with feeds in a simulated *in vitro* rumen environment; the control was constituted by the sterile WL growth medium. The digestibility of the Neutral Detergent Fibre content of the treated feeds (NDFd) was analysed and related to the level of *B. subtilis* enzymes supplied. NDFd is a parameter that defines the percentage of fibre that has been digested by ruminal microorganisms, which in turn is positively associated with forage quality (Weinberg et al., 2009; Nave et al., 2013). Analysing the pool of feed ingredients as a whole, a significant increment in NDFd was observed for both strains with respect to the control and between each other (p<0.001). In the samples treated with the WT *B. subtilis* strain, NDF digestibility attained to 45.6%, showing a 5% increase compared to the control (43.4% NDFd). Digestibility showed a remarkable 10% increment (47.8%) with respect to the control when the treatment was carried out with the higher amount of degradative enzymes secreted by the PB2OPT strain.

The NDFd data also revealed a significant interaction between Treatment x Feed (P=0.028). Particularly, the NDF content of maize silages (on average around 48% in our heterogeneous samples) was significantly more sensitive to enzymatic digestion and the degradation efficiency correlated with the amount of degradative enzymes released by the strains (PB2OPT>PB5700>Control) (Fig. 2); alfalfa hay showed a significant increment in NDFd only in the presence of the optimized strain, although a gradual trend was appreciable, whereas for meadow hay both strains equally contributed to the increase in NDFd as compared to the control (Fig. 2). Unsurprisingly, no effect of the bacterial treatment was observed on TMR forage, characterized by low NDF content (on average 34% in our samples) and already very digestible.

**Figure 2.**
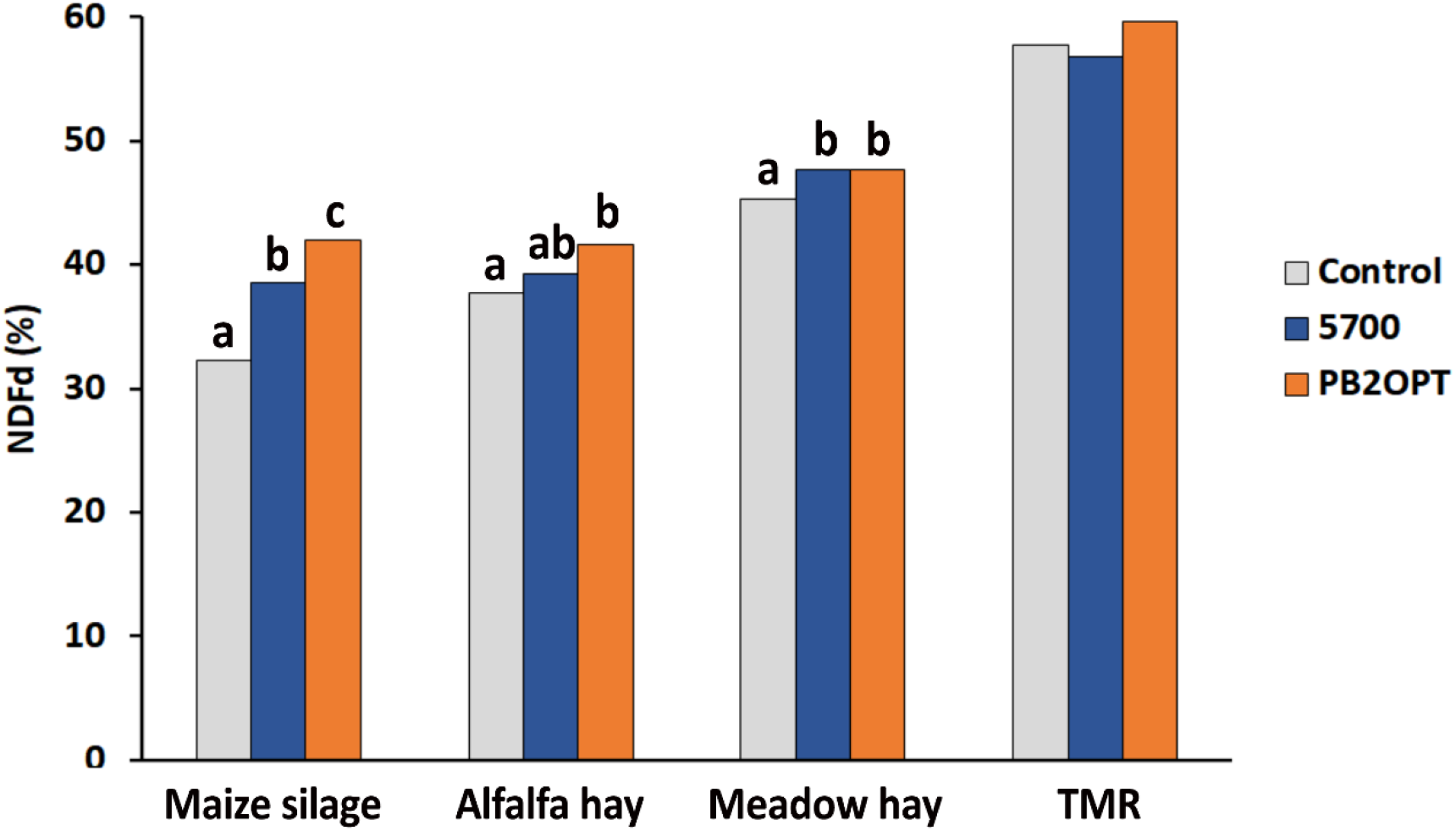
Fibre digestibility of forages. Fibre digestibility expressed as NDFd (%) in the four feed ingredients upon treatments: grey bars represent the negative control without bacteria; blue bars represent treatments with PB5700 strain; orange bars represent treatments with PB2OPT strain. Mean values in the same graph with different lettering are statistically different.

Indeed, the sensitivity to the treatment was inversely related to the intrinsic digestibility of the fibre, as represented by the NDFd of the untreated control samples, which in turn is linked to the chemical composition of the substrates. Maize silage, which is the most recalcitrant to ruminal activity among the substrates tested, benefited the most from the support of bacterial enzymes.

Although the increment given by the PB2OPT treatment compared to the WT is apparently not dramatic, accounting for 5% of NDFd, it is worth to recall that 1% increase in forage NDFd is associated with a 0.17 kg/d increase in DMI and 0.23 kg/d of milk produced (Oba and Allen, 1999; Kendall et al., 2009). This result demonstrates that *B. subtilis* cellulases and xylanases effectively contribute to increase fibre degradation in the rumen and can thus be responsible, at least in part, for the 14% increase in DMI and 6.94% higher milk production observed upon supplementation of *Bacillus subtilis* to lactating cows (Jia et al., 2022).

## 4. Conclusions

The effects of *Bacillus* spp. on *in vitro* digestibility of forages included in ruminant diets and its ability to increase in NDFd is not novel (Sun et al., 2013; Romero et al., 2015; Jia et al., 2022; Pan et al., 2022). The novelty of our work was the demonstration that the cellulases and xylanases secreted by this microorganism are effectively involved in the recovery of nutrients from fibrous material. This outcome was made possible by the application of two isogenic strains differing just for the production of these two specific enzymes. The comparison of the effect of their levels on feeds allowed to confidently ascribe to these two enzymes a relevant role in supporting the ruminal flora in the recovery of nutrients from fibrous material, excluding any confounding effect.

New strains, each engineered for the over-expression of a different set of enzymes, e.g., proteases, amylases, pullulanases, pectinases or others, might allow to verify whether each of them plays a role in the feed digestibility and, in the long run, might pave the way for developing the ideal strain to be used as feed additive.

Through the design of a convenient, sustainable, and simplified production process, feed additives, based on *Bacillus subtilis* raw cultures grown on an agricultural large waste biomass as rice straw, could be produced on site, and possibly be integrated in local biorefineries, creating a virtuous circular-economy process.

## Data availability

All data generated or analysed during this study are included in this published article.

## Acknowledgements

This research was supported by Fondazione Cariplo, Bando Economia Circolare: ricerca per un futuro sostenibile; grant # 2018-1011 and by the Italian Ministry of Education, University and Research (MIUR): Dipartimenti di Eccellenza Program (2018–2022) - Dept. of Biology and Biotechnology “L. Spallanzani”, University of Pavia.

## References

1. Adesogan, A.T., Arriola, K.G., Jiang, Y., Oyebade, A., Paula, E.M., Pech-Cervantes, A.A., Romero, J.J., Ferraretto, L.F., Vyas, D. 2019. Symposium review: Technologies for improving fibre utilization. J. Dairy Sci., 102: 5726–5755. DOI:10.3168/jds.2018-15334.

2. Belal, E.B. 2013. Bioethanol production from rice straw residues. Braz. J. Microbiol., 44: 225–34. DOI:10.1590/S1517-83822013000100033.

3. Bergstrand, M., Karlsson, M.O. 2009. Handling data below the limit of quantification in mixed effect models. AAPS J., 11: 371–80. DOI:10.1208/s12248-009-9112-5.

4. Bernardeau, M., Lehtinen, M. J., Forssten, S. D., Nurminen, P. 2017. Importance of the gastrointestinal life cycle of Bacillus for probiotic functionality. J. Food Sci. Technol., 54: 2570–2584. DOI:10.1007/s13197-017-2688-3.

5. Capper, J. L. and Bauman, D. E. 2013. The role of productivity in improving the environmental sustainability of ruminant production systems. Annu. Rev. Anim. Biosci., 1: 469–489. DOI:10.1146/annurev-animal-031412-103727.

6. Choonkham, W., Schonewille, J. T., Bernard, J. K., Suriyasathaporn, W. 2020. Effects of on-farm supplemental feeding of probiotic Bacillus subtilis on milk production in lactating dairy cows under tropical conditions. J. Anim. Feed Sci., 29: 199–205. DOI:10.22358/jafs/127692/2020

7. Danilova, I., Sharipova, M. 2020. The practical potential of bacilli and their enzymes for industrial production. Front. Microbiol., 11: 1782. DOI:10.3389/fmicb.2020.01782.

8. Donnelly, D. M., de Resende, L.C., Cook, D.E., Atalla, R.H., Combs, D.K. 2018. Technical note: A comparison of alkali treatment methods to improve neutral detergent fibre digestibility of corn stover. J. Dairy Sci., 101: 9058–9064. DOI:10.3168/jds.2017-14317.

9. Doria, E., Buonocore D., Marra A., Bontà V., Gazzola A., Dossena M., Verri M., Calvio C., 2022. Bacterial-Assisted Extraction of Bioactive Compounds from Cauliflower. Plants 11: 816. DOI:10.3390/plants11060816.

10. Earl, A. M., Losick, R., Kolter, R. 2008. Ecology and genomics of *Bacillus subtilis*. Trends Microbiol. 16: 269–275. DOI:10.1016/j.tim.2008.03.004.

11. EFSA BIOHAZ Panel. 2022. Updated list of QPS-recommended biological agents for safety risk assessments carried out by EFSA [Data set]. Zenodo. DOI:10.5281/zenodo.6902983.

12. EFSA BIOHAZ Panel. 2023a. Microbiological agents as notified to EFSA [Data set]. Zenodo. DOI:10.5281/zenodo.7554134.

13. EFSA BIOHAZ Pane. 2023b. Scientific Opinion on the update of the list of qualified presumption of safety (QPS) recommended microorganisms intentionally added to food or feed as notified to EFSA. EFSA J., 21: 7747. DOI:10.2903/j.efsa.2023.7747

14. Ermoli, F., Bontà, V., Vitali, G., Calvio, C. 2021. SwrA as global modulator of the two-component system DegSU in *Bacillus subtilis*. Microbiol. Res., 172: 103877. DOI: 10.1016/j.resmic.2021.103877.

15. Ferreira, R. G., Azzoni, A. R., Freitas, S. 2021. On the production cost of lignocellulose-degrading enzymes. Biofuels, Bioprod. Bior., 15: 85–99. DOI:10.1002/bbb.2142.

16. Himmel, M. E., Xu, Q., Luo, Y., Ding, S. Y., Lamed, R., Bayer, E. A. 2010. Microbial enzyme systems for biomass conversion: emerging paradigms. Biofuels, 1: 323–341. DOI:10.4155/bfs.09.25.

17. Jia, P., et al. 2022. Diets supplementation with Bacillus subtilis and Macleaya cordata extract improve production performance and the metabolism of energy and nitrogen, while reduce enteric methane emissions in dairy cows. Anim. Feed Sci. Technol., 294: 115481. DOI:10.1016/j.anifeedsci.2022.115481.

18. Kendall, C., Leonardi, C., Hoffman, P. C., Combs, D. K. 2009. Intake and milk production of cows fed diets that differed in dietary neutral detergent fibre and neutral detergent fibre digestibility. J. Dairy Sci., 92: 313–323. DΘI:10.3168/jds.2008-1482.

19. Krishania, M., Kumar, V., Singh Sangwan, R. 2018. Integrated approach for extraction of xylose, cellulose, lignin and silica from rice straw. Bioresour. Technol. Rep., 1: 89–93. DOI:10.1016/j.biteb.2018.01.001.

20. Lee, N.K., Kim, W.S. Paik, H.D. 2019. Bacillus strains as human probiotics: characterization, safety, microbiome, and probiotic carrier. Food Sci. Biotechnol., 28: 1297–1305. DOI: 10.1007/s10068-019-00691-9

21. Lombard, V., Golaconda Ramulu, H., Drula, E., Coutinho, P. M., Henrissat, B. 2016. The carbohydrate-active enzymes database (CAZy) in 2013. Nucleic Acids Res., 42: 490–495. DOI:10.1093/nar/gkt1178.

22. McIntosh, S., Vancov, T. 2011. Optimization of dilute alkaline pretreatment for enzymatic saccharification of wheat straw. Biomass Bioenerg., 35: 3094–3103. DOI:10.1016/j.biombioe.2011.04.018.

23. Mertens, D.R. 2002. Gravimetric determination of amylase-treated neutral detergent fibre in feeds with refluxing in beakers or crucibles: collaborative study. J. AOAC Int. 85:1217–1240.

24. Mor, P., et al. 2018. Effect of ammonia fibre expansion on the available energy content of wheat straw fed to lactating cattle and buffalo in India. J. Dairy Sci., 101: 7990–8003. DOI:10.3168/jds.2018-14584.

25. Mun, D., et al. 2021. Effects of *Bacillus-based* probiotics on growth performance, nutrient digestibility, and intestinal health of weaned pigs. J Anim Sci Technol., 63:1314–1327. DOI:10.5187/jast.2021.e109.

26. Nave, R. L. G., Sulc, R. M., Barker, D. J. 2013. Relationships of forage nutritive value to cool-season grass canopy characteristics. Crop Sci., 53: 341–348. DOI:10.2135/cropsci2012.04.0236.

27. Oba, M. and Allen, M.S. 1999. Evaluation of the Importance of the Digestibility of Neutral Detergent Fibre from Forage: Effects on Dry Matter Intake and Milk Yield of Dairy Cows. J. Dairy Sci., 82: 589–596. DOI:10.3168/jds.S0022-0302(99)75271-9.

28. Pan, L., Harper, K., Queiroz, O., Copani, G., Cappellozza, B.I. 2022. Effects of a Bacillus-based direct-fed microbial on in vitro nutrient digestibility of forage and high-starch concentrate substrates. Transl. Anim. Sci., 6. DOI:10.1093/tas/txac067.

29. Park, J., Seyama, T., Shiroma, R., Ike, M., Srichwong, S., Nagata K., Arai-Sanoh, Y., Kondo, M., Tokuyasu, K. 2009. Efficient Recovery of Glucose and Fructose via Enzymatic Saccharification of Rice Straw with Soft Carbohydrates. Biosci. Biotechnol. Biochem., 73:1072–1077. DOI:10.1271/bbb.80840.

30. Romero, J.J., Zarate, M.A., Arriola, K.G., Gonzalez C.F., Silva-Sanchez, C., Staples, C.R., Adesogan, A.T. 2015. Screening exogenous fibrolytic enzyme preparations for improved in vitro digestibility of bermudagrass haylage. J. Dairy Sci., 98: 2555–67. DOI:10.3168/jds.2014-8059.

31. Schallmey, M., Singh, A., Ward, O. P. 2004. Developments in the use of Bacillus species for industrial production. Can. J. Microbiol., 50: 1–17. DOI:10.1139/w03-076.

32. Smith, J. L., Goldberg, J. M., Grossman, A. D. 2014. Complete Genome Sequences of *Bacillus* subtilis subsp. subtilis Laboratory Strains JH642 (AG174) and AG1839. Genome Announc., 2. DOI:10.1128/genomeA.00663-14.

33. Souza, V.L., et al. 2017. Lactation performance and diet digestibility of dairy cows in response to the supplementation of Bacillus subtilis spores. Livest. Sci., 200: 35–39. DOI:10.1016/j.livsci.2017.03.023.

34. Spanghero, M., et al. 2019. Rumen Inoculum Collected from Cows at Slaughter or from a Continuous Fermenter and Preserved in Warm, Refrigerated, Chilled or Freeze-Dried Environments for In Vitro Tests. Animals (Basel), 9: 815. DOI:10.3390/ani9100815.

35. Srivatsan, A., et al. 2008. High-precision, whole-genome sequencing of laboratory strains facilitates genetic studies. PLoS Genet., 4. DOI:10.1371/journal.pgen.1000139.

36. Su, Y., Liu, C., Fang, H., Zhang, D. 2020. Bacillus subtilis: A universal cell factory for industry, agriculture, biomaterials and medicine. Microb. Cell Fact., 19: 1–12. DOI:10.1186/s12934-020-01436-8.

37. Sun, P., Wang, J. Q., Zhang, H. T. 2011. Effects of supplementation of Bacillus subtilis natto Na and N1 strains on rumen development in dairy calves. Anim. Feed Sci. Tech., 64: 154–160. DOI:10.1016/j.anifeedsci.2011.01.003.

38. Sun, P., Wang, J. Q., Deng, L. F. 2013. Effects of Bacillus subtilis natto on milk production, rumen fermentation and ruminal microbiome of dairy cows. Animal, 7: 216–222. DOI:10.1017/S1751731112001188.

39. Sun, P., Li, J., Bu, D., Nan, X., Du H. 2016. Effects of *Bacillus subtilis natto* and Different Components in Culture on Rumen Fermentation and Rumen Functional Bacteria In Vitro. Curr. Microbiol., 72: 589–595. DOI:10.1007/s00284-016-0986-z.

40. van Dijl, J. and Hecker, M. 2013. *Bacillus subtilis:* from soil bacterium to super-secreting cell factory. Microb.Cell. Fact., 12. DOI:10.1186/1475-2859-12-3.

41. Wang, Q., et al. 2022. *Bacillus subtilis* Produces Amino Acids to Stimulate Protein Synthesis in Ruminal Tissue Explants *via* the Phosphatidylinositol-4,5-Bisphosphate 3-Kinase Catalytic Subunit Beta-Serine/Threonine Kinase-Mammalian Target of Rapamycin Complex 1 Pathway. Front. Vet. Sci., 27: 852321. DOI:10.3389/fvets.2022.852321.

42. Weinberg Z.G., Chen Y., Solomon R. 2009. The quality of commercial wheat silages in Israel, J. Dairy Sci., 92: 638–644.DOI:10.3168/jds.2008-1120.

43. Westers, L., Westers, H., Quax, W. J. 2004. *Bacillus subtilis* as cell factory for pharmaceutical proteins: A biotechnological approach to optimize the host organism. Biochim.Biophys. Acta - Mol. Cell Res., 1694: 299–310. DOI:10.1016/j.bbamcr.2004.02.011.

